## Supplemental Figures for "Structural insights into the mechanism of rhodopsin phosphodiesterase"

#### Extended Data Figures

##### Extended Data Fig. 1. Purification and crystallization of Rh-PDE-TMD.

**a**, Gel filtration chromatogram of the purified Rh-PDE-TMD. Fractions with dashed lines were collected for the following crystallization. **b**, Crystals of Rh-PDE-TMD. **c,** Structural comparison between mol A and B. Mol A and B are colored rainbow and light orange, respectively.

##### Extended Data Fig. 2. Overall structural comparison with ChR2 and HeR.

Overall structures of Rh-PDE (left), ChR2 (middle) and HeR (right), viewed from the membrane plane (upper) and extracellular side (lower). Monoolein molecules are colored yellow.

##### Extended Data Fig. 3. Sequence alignment with Rh-PDE homologs.

Amino acid sequence alignment of *Sr*Rh-PDE and other Rh-PDE homologs, from *Microstomoeca roanoke*, *Choanoeca perplexa*, and *Choanoeca flexa*. The star indicates the conserved lysine residues in TM7.

##### Extended Data Fig. 4. Surface residues and TM0 deletion mutants.

**a,** Conservation of the surface residues of the Rh-PDE structure. The sequence conservation among seven Rh-PDE homologs was calculated using the ConSurf server (http://consurf.tau.ac.il) and is coloured from cyan (low) to maroon (high). **b,**

Photoactivities of truncated mutants with cAMP (left) and cGMP (right).

#### Extended Data Fig. 5. Purification and crystallization of Rh-PDE TMD-Linker.

**a**, Gel filtration chromatogram of the purified Rh-PDE TMD-Linker. Fractions with dashed lines were collected for the following crystallization. **b**, Crystals of Rh-PDE TMD-Linker. **c,** Crystal packing of Rh-PDE TMD-Linker. **d,** Structural comparison between Rh-PDE TMD and TMD-Linker. TMD and TMD-Linker are colored orange and rainbow, respectively. **e,** Structural comparison between Rh-PDE TMD mol A and TMD-Linker mol B. TMD and TMD-Linker are colored light and dark orange, respectively. **f,** 2Fo-Fc map of the linker region of Rh-PDE TMD-Linker, contoured at 1.0σ.

#### Extended Data Fig. 6. Structural comparison of PDE domains.

**a,** Structural comparison with the previous reported Rh-PDE PDE domain (PDB ID: 5VYD). Our structure and the previous structure are colored orange and gray, respectively. N-terminal residues of each structure are shown as spheres. **b,** Structural comparison with the PDE9 catalytic domain (PDB ID: 2HD1). Our structure and the PDE9 structure are colored orange and blue, respectively. N-terminal residues of each structure are shown as spheres.

#### Extended Data Fig. 7. Structural comparison of the linker of aMD simulations.

**a-c,** Structural comparison between the linker model shown in Fig. 4d and the final structures of aMDd (**a**), aMDT (**b**) and aMDdual (**c**). The linker model and each aMD structure are colored gray and rainbow, respectively. Cα atoms of I311 and M386 are shown as spheres.

**Extended Data Fig. 8. HS-AFM.**

Simulation images of PDE dimers, from three viewpoints.

**Extended Data Fig. 9. Sequence alignment with rhodopsin enzymes.**

Amino acid sequence alignment of *Sr*Rh-PDE TMD and corresponding regions of rhodopsin enzymes, from *Blastocladiella emersonii* rhodopsin-guanylyl cyclase 1 (*Be*Rh-GC1), *Catenaria anguillulae* rhodopsin-guanylyl cyclase (*Ca*Rh-GC), *Chlamydomonas reinhardtii* two-component cyclase opsin 1 (*Cr*2c-Cyclop1) and *Volvox carteri* two-component cyclase opsin 1 (*Vr*2c-Cyclop1). The star indicates the conserved lysine residues in TM7.

**Extended Data Table 1. Data collection and refinement statistics.**

| | TMD | TMD-Linker | Linker-PDE |
|---|---|---|---|
| **Data collection** | | | |
| Space group | *P*222 | *I*222 | *C*2 |
| Cell dimensions | | | |
| *a*, *b*, *c* (Å) | 65.55, 74.14, 117.38 | 76.30, 136.53, 206.54 | 117.2, 67.54, 56.89 |
| α, β, γ (°) | 90, 90, 90 | 90, 90, 90 | 90, 110.817, 90 |
| Resolution (Å)* | 49.11 - 2.6 | 49.40 - 3.5 | 47.53 - 2.1 |
| | (2.76 - 2.6) | (3.71 - 3.5) | (2.18 - 2.1) |
| $R_{meas}$* | 0.845 (13.2) | 0.673 (2.868) | 0.162 (0.798) |
| $<I/\sigma(I)>$ * | 7.13 (0.76) | 3.90 (1.04) | 11.4 (2.5) |
| $CC_{1/2}$* | 0.996 (0.609) | 0.934 (0.393) | 0.992 (0.660) |
| Completeness (%)* | 99.9 (100) | 98.1 (98.5) | 90.7 (81.7) |
| Redundancy* | 62.4 (57.0) | 6.55 (6.18) | 11.6 (8.7) |
| **Refinement** | | | |

| | | | |
|---|---|---|---|
| Resolution (Å) | 49.11 - 2.6 | 49.40 - 3.5 | 47.53 - 2.1 |
| No. reflections | 18,193 | 13,712 | 22,066 |
| $R_{work}$ / $R_{free}$ | 0.2458 / 0.2950 | 0.2663 / 0.3144 | 0.1883 / 0.2397 |
| No. atoms | 4,402 | 4,611 | 2,814 |
| Protein | 4,052 | 4,485 | 2,600 |
| Ligand/ion | 334 | 126 | 38 |
| Water | 16 | 0 | 176 |
| Averaged *B*-factors ($Å^2$) | 40.43 | 33.37 | 27.78 |
| Protein | 39.35 | 33.48 | 26.99 |
| Ligand/ion | 54.37 | 29.29 | 42.23 |
| Water | 23.06 | | 36.21 |
| R.m.s. deviations from ideal | | | |
| Bond lengths (Å) | 0.016 | 0.003 | 0.004 |
| Bond angles (°) | 1.81 | 0.61 | 0.88 |
| Ramachandran plot | | | |
| Favored (%) | 97.32 | 95.55 | 99.08 |
| Allowed (%) | 2.49 | 4.11 | 0.92 |
| Outlier (%) | 0.19 | 0.34 | 0 |

*Values in parentheses are for highest-resolution shell.

#### Supplementary Video 1. HS-AFM movie of the full-length Rh-PDE reconstituted in a lipid bilayer.

Scan area: 60 × 35 $nm^2$ (140 × 84 pixels). Imaging rate: 0.1 s/frame.

**Supplementary Video 2. HS-AFM movie of the full-length Rh-PDE reconstituted in a lipid bilayer.**

Scan area: 41 × 28 $nm^2$ (96 × 67 pixels). Imaging rate: 0.1 s/frame.

**Supplementary Video 3. HS-AFM movie of the full-length Rh-PDE reconstituted in a lipid bilayer.**

Scan area: 65 × 52 $nm^2$ (140 × 96 pixels). Imaging rate: 0.1 s/frame.

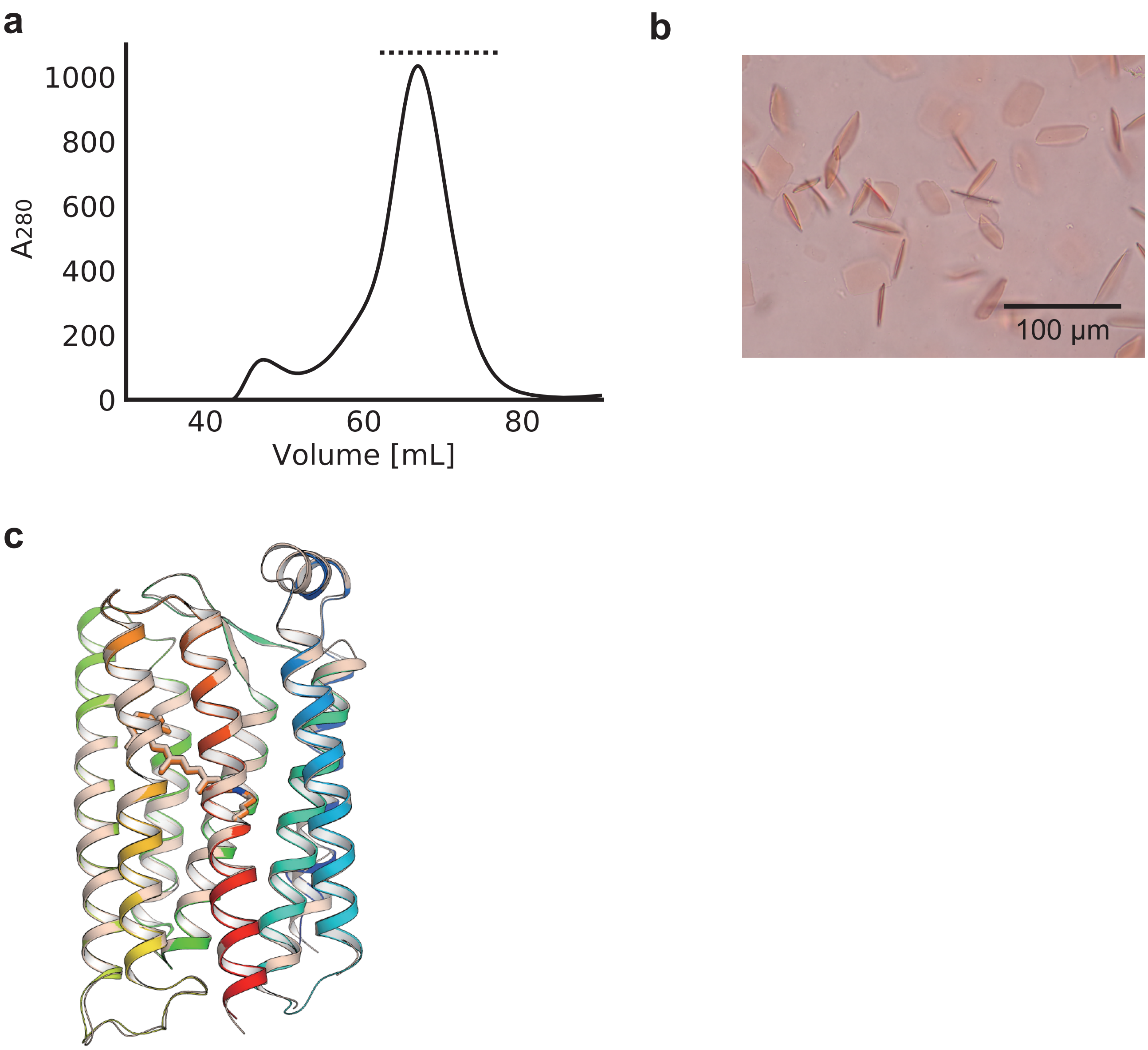


### Extended Data Fig. 1

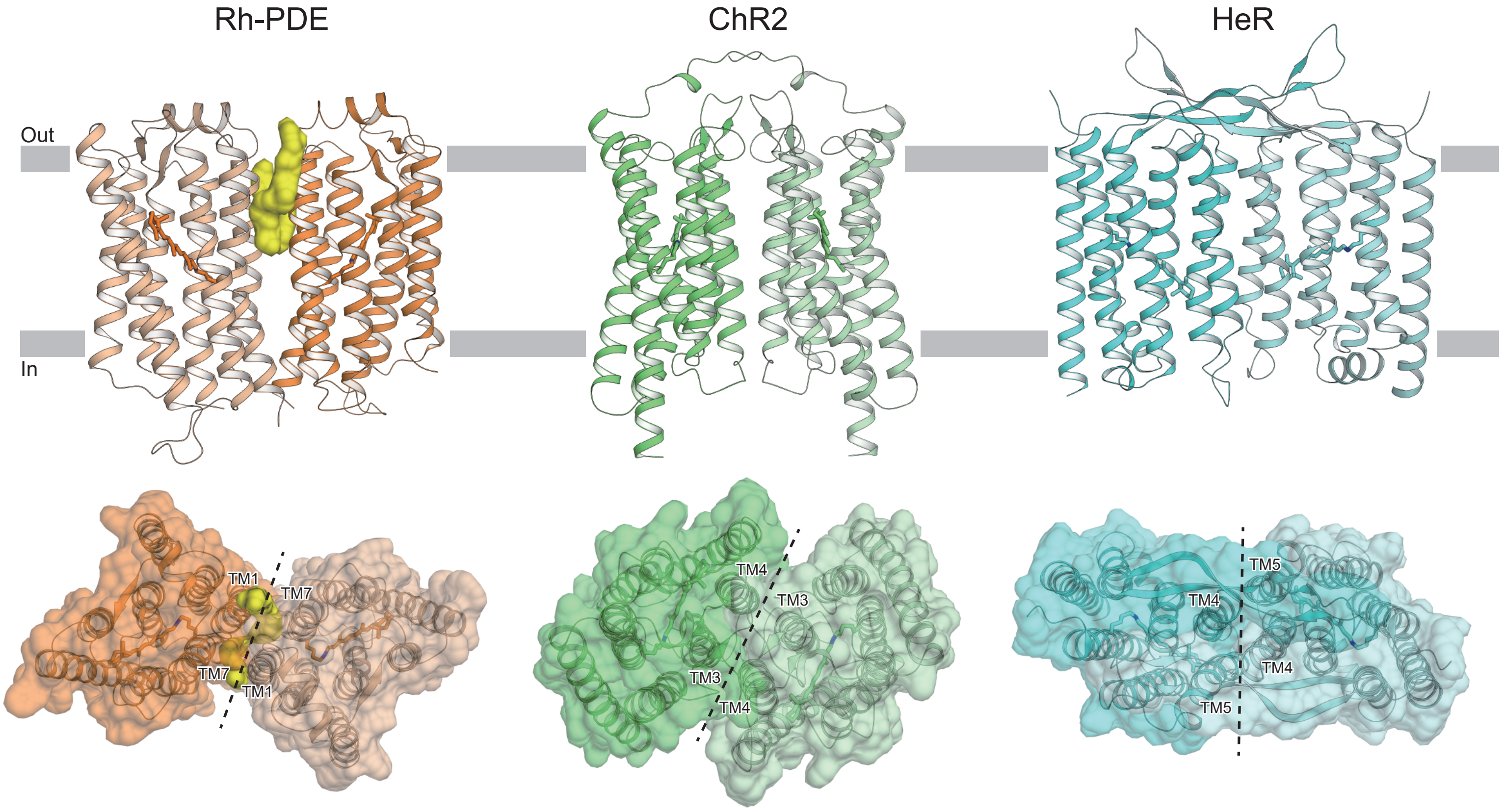


### Extended Data Fig. 2

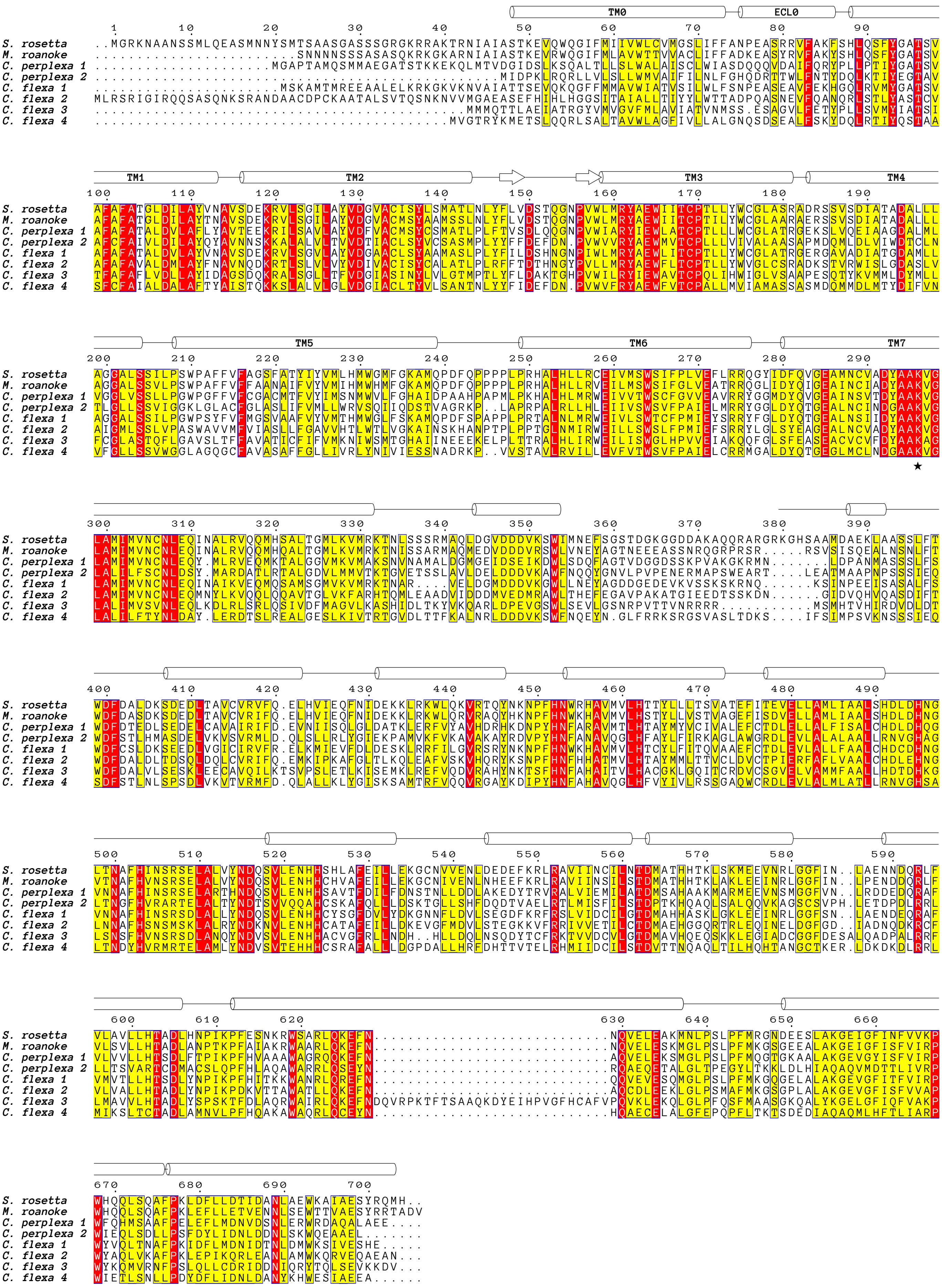


### Extended Data Fig. 3

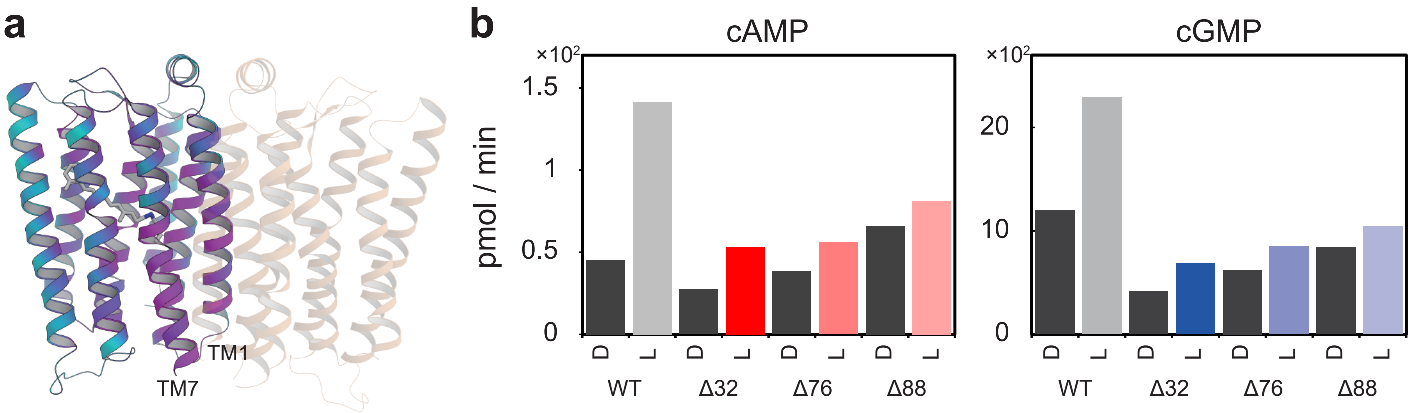


### Extended Data Fig. 4

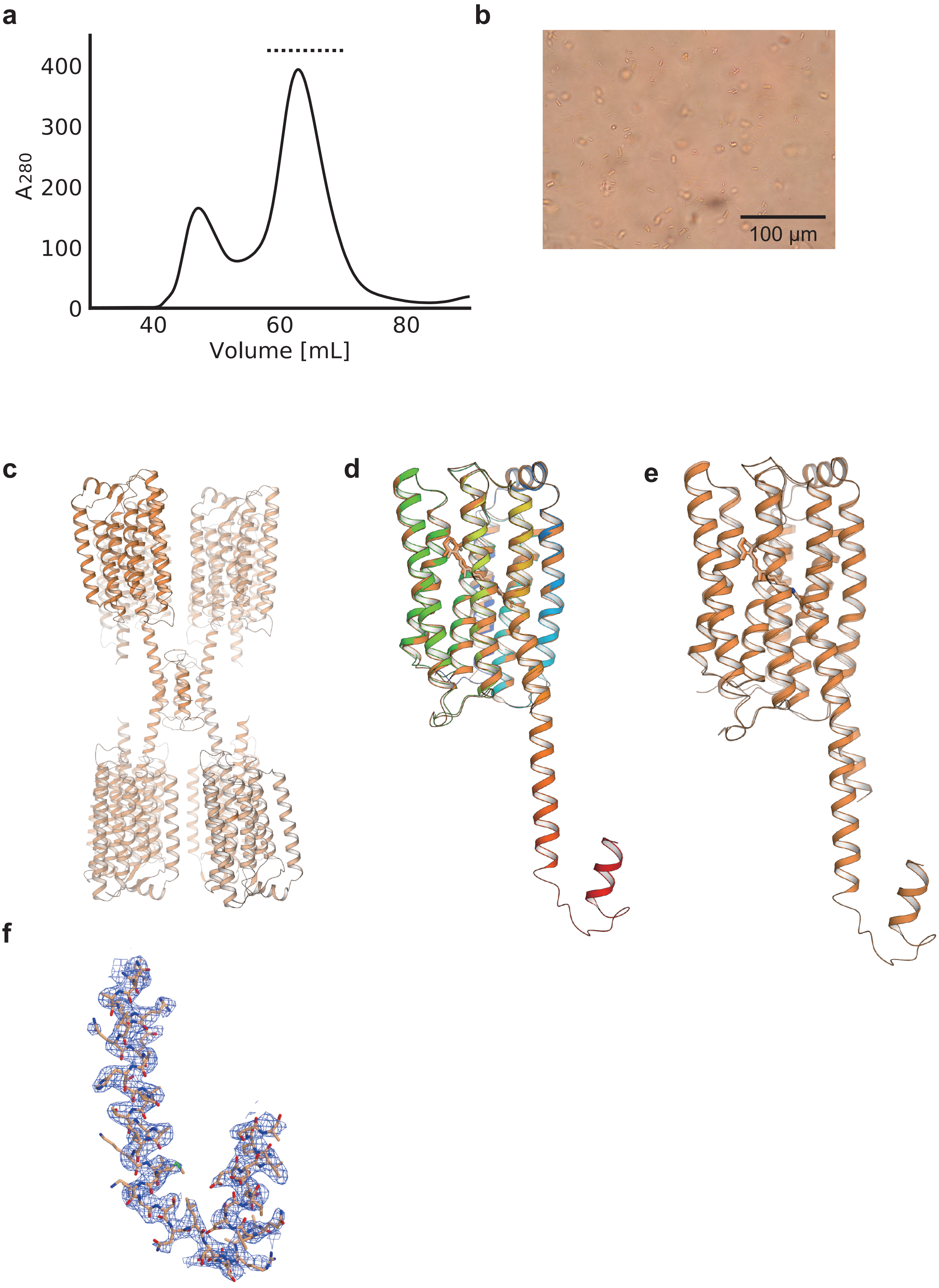


### Extended Data Fig. 5

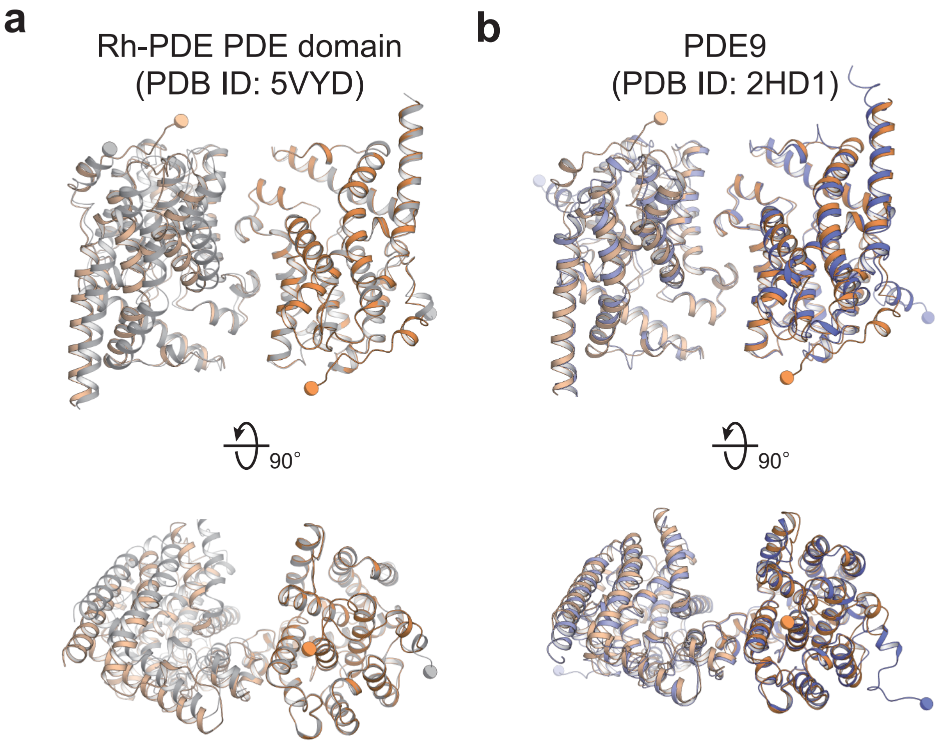


### Extended Data Fig. 6

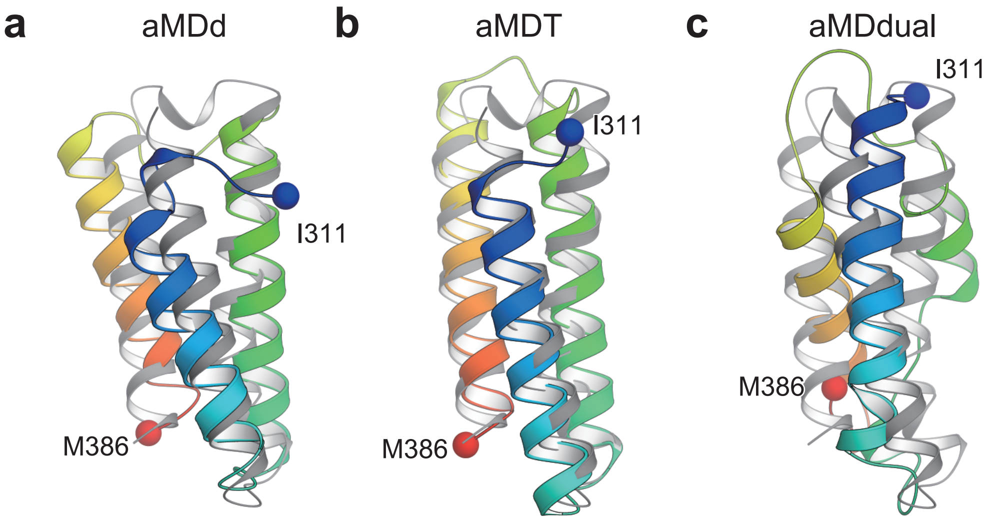


### Extended Data Fig. 7

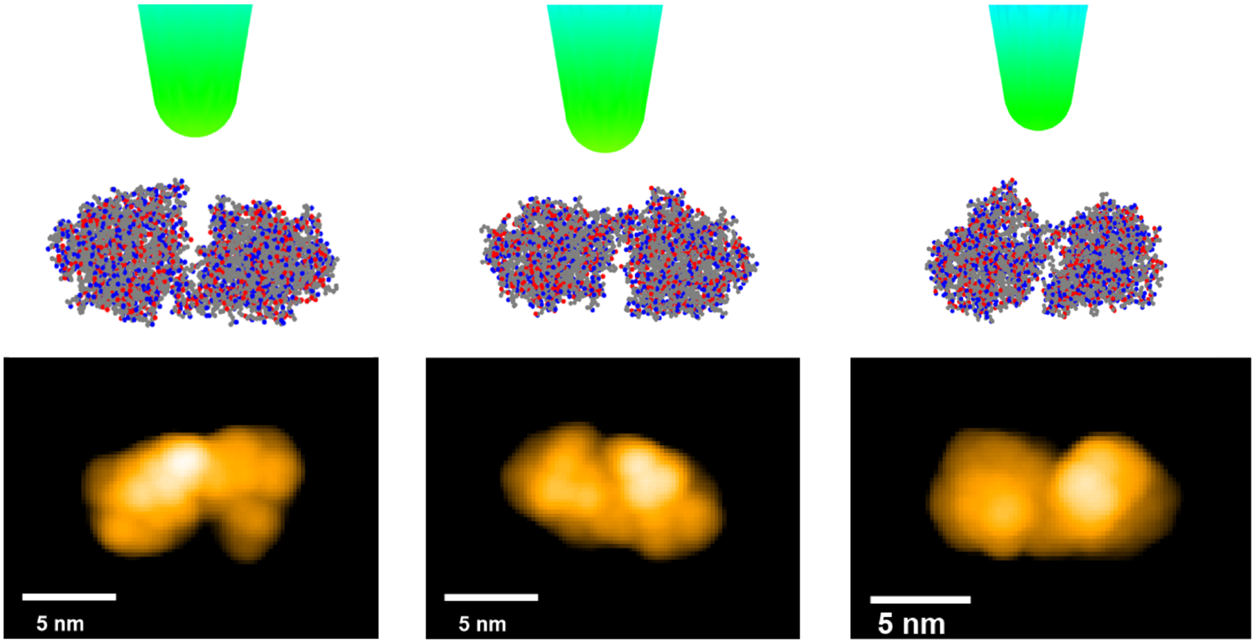


### Extended Data Fig. 8

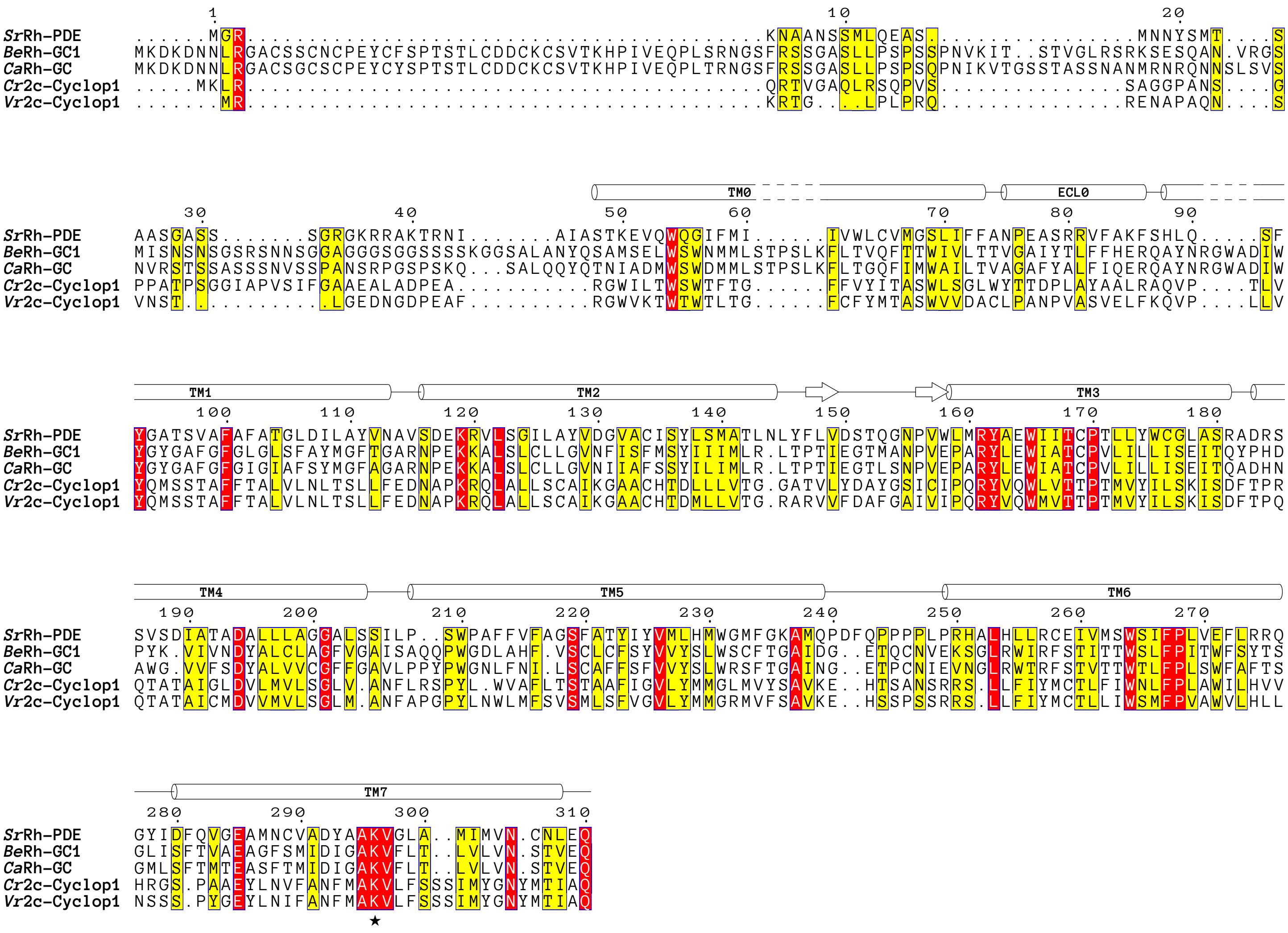


### Extended Data Fig. 9
